# Forest tree species composition and abiotic site conditions drive soil fungal communities and functional groups

**DOI:** 10.1101/2021.07.21.453256

**Authors:** Likulunga Emmanuel Likulunga, Carmen Alicia Rivera Pérez, Dominik Schneider, Rolf Daniel, Andrea Polle

**Affiliations:** Forest Botany and Tree Physiology, University of Goettingen, Büsgenweg 2, 37077 Göttingen, Germany; Biological Sciences Department, University of Zambia, Great East Road Campus, 32379 Lusaka, Zambia; Genomic and Applied Microbiology and Göttingen Genomics Laboratory, University of Göttingen, 37077 Göttingen, Germany

**Author notes:** Corresponding author: E-mail address (A. Polle), Address: Forest Botany and Tree Physiology, Faculty of Forest Sciences, University of Göttingen, Büsgenweg 2, 37077, Göttingen, Germany.

**Keywords:** Fungal communities, pure and mixed tree stands, symbiotrophs, saprotrophs, soil properties, ITS2

## Abstract

Soil fungi, especially the functional guilds of saprotrophic and mycorrhizal fungi, play a central role in ecosystem processes by degrading litter, mining for mineral nutrients and linking above- and belowground nutrient fluxes. Fungal community structures are influenced by abiotic habitat filters and management decisions such as tree species selection. Yet, the implications of the enrichment of temperate forests consisting of tree species in their natural range with non-native tree species on soil fungal diversity and their functional groups are unknown. Here, we studied fungal communities in 40 plots located in two regions differing in site conditions (nutrient content and soil moisture) in forests composed of European beech, Norway spruce and Douglas-fir (non-native) and mixtures of beech with either spruce or Douglas-fir. We hypothesized that fungal community structures are driven by soil resources and tree species composition, generally resulting in higher fungal diversity in mixed than in mono-specific forests. We further hypothesized that Douglas-fir has a negative effect on ectomycorrhizal fungal species richness compared to native species, whereas saprotrophic fungal richness is unaffected. We found strong separation of fungal communities between nutrient-rich and nutrient-poor sites and taxonomic divergence between beech and conifer fungal communities and an intermediate pattern in mixed forests. Mycorrhizal species richness did not vary with forest type, but the relative abundance of mycorrhizal species was lower in Douglas-fir and in mixed beech-Douglas-fir forests than in spruce or beech-spruce mixture. Conifer forests contained higher relative abundances of saprotrophic fungi than mono-specific beech forests. Among 16 abundant fungal orders in soil, two containing saprotrophic fungi (Tremellales, Hymenochaetales) were enriched in conifer forests, regardless of site conditions and tree species mixture. The other fungal orders, including those dominated by mycorrhizal fungi (Russulales, Boletales, Atheliales, Cantharellales) showed variable patterns depending on site conditions and tree species. In conclusion, Douglas-fir mono-specific or mixed forests show no loss of fungal species richness, but a shift in functional composition towards saprotrophic fungi.

## 1. Introduction

Ecosystem processes such as nutrient cycling, decomposition, nutrient acquisition and protection against plant diseases are mediated by soil microbes (Bardgett and Wardle, 2010). In temperate and boreal forests, soil fungi contribute substantially to these functions (Baldrian, 2017; Brundrett and Tedersoo, 2018). Based on carbon acquisition strategies, soil fungi can be divided into saprotrophic, symbiotrophic and pathotrophic fungal categories (Schmit and Mueller, 2007; Nguyen et al., 2016). Saprotrophic fungi are the main decomposers of litter and wood (Rayner and Boddy, 1988), while symbiotrophic fungi form mutualistic associations with roots facilitating plant nutrient uptake in exchange for carbohydrates (Smith and Read, 1997). Since fungi play a fundamental role in ecosystem processes, it is important to understand how forest management affects the composition of fungal guilds.

In large parts of Europe, European beech (*Fagus sylvatica*) and Norway spruce (*Picea abies*) are ecologically and economically important tree species (Leuschner et al., 2006; Leuschner and Ellenberg, 2017). Due to climate change and calamities, which jeopardize beech and spruce forests (Schlyter et al., 2006; Gessler et al., 2007; Bolte et al., 2010), there is the need to introduce tree species that resist these changes and show compatibility with beneficial soil microbes. Douglas-fir (*Pseudotsuga menziesii*), which is native to North America (Hermann, 1987), is one of the long-term introduced species in Europe (Essl, 2005). Douglas-fir exhibits desirable growth characteristics (Sicard et al., 2006; von Lüpke, 2009; Isaac-Renton et al., 2014) and the ability to thrive well with other broadleaved and coniferous tree species (Rothe and Binkley, 2001). These characteristics make Douglas-fir a potential candidate for European forestry. However, little is known about the influence of Douglas-fir on belowground fungal communities. Previous studies conducted with Douglas-fir growing outside its natural range focused on the ability to form mycorrhizal associations with resident fungal species (Schmid et al., 2014; Moeller et al., 2015; Parlade et al., 1995; Dučić et al., 2009). Information on the impact of Douglas-fir on soil fungal diversity and composition under varying site conditions and soil properties in temperate European forests is lacking.

In general, the taxonomic and functional composition of soil fungi are driven by abiotic and biotic environmental factors such as soil pH, moisture, C/N ratio, and the composition of the vegetation (Kivlin et al., 2014; Wubet et al., 2012; Bahnmann et al., 2018). The latter has strong influence on litter input, shading, rain interception, transpiration and root exudation (Stoutjesdijk and Barkman, 1992; Augusto et al., 2002). Consequently, traits of forest species influence habitat properties (Bahnmann et al., 2018; Põlme et al., 2018; Urbanová et al., 2015). For example, litter of conifers (spruce and Douglas-fir) contains lower nitrogen (N) and cation contents than that of beech trees (Kubartová et al., 2009). In conifer stands of Douglas-fir or spruce, soil organic carbon and N stocks are higher than in beech stands (Cremer et al., 2016; Cremer and Prietzel, 2017; Dawud et al., 2017). Douglas-fir and beech forest soils exhibit higher exchangeable calcium and magnesium concentrations than soil in spruce or mixed beech-spruce stands (Foltran et al., 2020). Changes in soil resources impact saprotrophic and symbiotrophic fungal guilds (Lindahl et al., 2007). Recent studies suggest that the phylogenetic composition of both guilds is responding to different ecological factors, leading to divergent communities in response to environmental changes (Schröter et al. 2019; Nguyen et al. 2020; Ballauff et al., 2021). Therefore, we anticipated that the introduction of Douglas-fir into European forests has profound effects on the functional and taxonomic composition of soil fungal communities.

Here, we investigated whether Douglas-fir cultivation, pure or in mixture with beech affects soil fungal richness, diversity and community composition in comparison with spruce or beech. We conducted our study in forests composed of either pure stands of beech, spruce, and Douglas-fir or of beech-conifer mixtures (beech-spruce, beech- Douglas-fir) stocking on nutrient-rich silt/clay containing soils as well as on nutrient-poor, sandy soils (Foltran et al. 2020). We used this experimental design to test the following hypotheses: i) Douglas-fir results in reduced soil fungal richness compared with beech or spruce because the fungal community is less adapted to an introduced than to native tree species. ii) Each stand type is characterized by a distinct soil fungal community composition because of the influence of abiotic habitat filters and of tree identity. Therefore, mixed stands contain highest fungal richness and diversity. Alternatively, we expected that the soil mycobiome is adapted to a wide range of conditions and therefore, no differences in fungal species richness occur. iii) Soils under conifers exhibit higher saprotrophic fungal richness and diversity than those under beech because lower litter quality of conifers favors decomposer communities (Cornelissen et al., 2001; Kubartová et al., 2009). iv) We reasoned that more closely related fungi will show similar responses to changes in habitat conditions than phylogenetically distant fungi. Therefore, we hypothesized that a specific impact of Douglas-fir is reflected by shifts in fungal orders compared with beech or spruce and occurs irrespective of abiotic site conditions.

## 2. Material and Methods

### 2.1. Study sites

The study was conducted in eight study sites (Harz, Dassel, Winnefeld, Nienover, Nienburg, Unterlüss, Göhrde II and Göhrde I) located in Germany. In each of the study sites, five plots with an area of 2500 m^2^ were established in the different forest types, which were composed of either European beech (B) (*Fagus sylvatica*), Norway spruce (S) (*Picea abies*), Douglas-fir (D) (*Pseudotsuga menziesii*), the mixture of European beech with Norway spruce (BS) or the mixture of European beech with Douglas-fir (BD). The stand age varied from 41 to 129 years. All plots were limed with approximately 1.9 t ha^-1^ CaCO_3_ and 0.8 t ha^-1^ MgCO_3_ applied in two doses (one at end of the 1980’s and in the first decade of the 2000’s) and seven plots (Unterlüss: D, BD, B and BS; Göhrde I: D, BD and S) were fertilized with P_2_O_5_ (0.24 t ha^-1^ approximately 30 years ago). The stand characteristics and climatic conditions are available in PANGEA (Ammer et al., 2020). Briefly, the sites Harz (51°46′12" N; 10°23′49" E), Dassel (51°42′33" N; 9°43′14" E), Winnefeld (51°39′52" N; 9°34′19" E) and Nienover (51°41′54" N; 9°31′47" E) are at elevations ranging from 282 m to 520 m above sea level (a.s.l) exhibiting high annual precipitation (869 to 1029 mm), while the sites Nienburg (52°37′14" N; 9°16′52" E), Unterlüss (52°50′12" N; 10°19′51" E), Göhrde I (53°12′4" N; 10°48′3" E) and Göhrde II (53°7′35" N; 10°48′48" E) are on lower elevation levels (84m to 164 m a.s.l) with lower precipitation (672 to 733 mm). The mean annual temperature across all study sites ranged from 7.63 to 9.70 ℃. The proportion of conifers varied from 9.4 to 58.5 % in the beech-conifer mixed stands (Ammer et al., 2020). The basal stem area ranged from 24.3 to 56.3 m^2^ ha^-1^ across all study sites (Ammer et al., 2020). The study sites exhibit differences in soil properties: Harz soil has a high clay content, Dassel, Winnefeld and Nienover exhibit high silt contents with low clay contents and the sites Nienburg, Unterlüss, Göhrde I and II have high sand contents with very low clay contents (Foltran et al., 2020).

Each plot was divided into quadrants (North East, South East, South West and North West). The North West quadrant of the plot was not used for destructive sampling. In each of the other three subplots, five soil cores (8 cm diameter x 10 cm depth) were taken in late fall (November/December, 2017), after removal of coarse litter in non- decomposing state (L layer). The five soil cores per subplot were pooled to yield one sample. Thereby, we collected three replicates per forest in each study site. Soil samples were immediately transported in cooling boxes with ice to the laboratory. The soil was sieved (4 mm mesh size) and divided in three aliquots: fresh aliquots frozen at - 20 ℃ (soil fungal composition analyses), aliquots dried at 40 ℃ (soil chemistry), and aliquots dried at 105 ℃ (soil moisture analyses).

### 2.2. Soil chemistry

To determine soil pH, 25 ml of water was added to 10 g of oven-dried soil (40 ℃), followed by the addition of a spatula tip of KCl. The suspension was shaken for 2h. After sedimentation of particles, the pH was measured (WTW pH meter 538, Wissenschaftlich-Technische-Werkstätten, Weilheim, Germany). To calculate means, the pH values were converted to proton (H^+^) concentrations, from which means (from three measurements) were calculated and back-transformed to pH.

Relative soil moisture was determined by drying soil samples at 105 ℃ for 72 hours and calculated as:

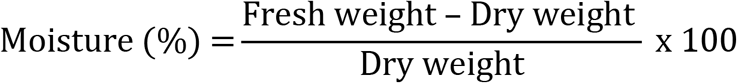

To measure soil C and N contents, oven-dried soil samples (40 ℃) were milled in a ball mill (MN400, Retsch GmbH, Haan, Germany). About 20 mg of milled soil samples were weighed into 4 x 4 x 11 mm tin capsules (IVA Analysentechnik, Meerbusch, Germany) on a microbalance (Model: Cubis MSA 2.7S-000-DM, Sartorius, Göttingen, Germany) and analyzed in a CN analyzer (vario MICRO cube CN analyzer, Elementar Analysensysteme, GmbH, Langenselbold Germany). We used acetanilide (10.36% N, 71.09% C) as the standard.

Further nutrient elements were extracted from oven-dried, milled soil samples by microwave digestion and determined by inductively coupled plasma–optical emission spectroscopy (ICP-OES) as follows: ultra-pure water (arium® pro, Sartorius Lab Instruments GmbH & Co. KG, Duderstadt, Germany) was added to 2 ml of 65 % HNO_3_ up to the final volume of 25 ml followed by addition of a defined weight (about 50 mg) of the milled soil sample. The sample was extracted by digestion in a microwave (Etho.start, Mikrowellen-Labor-Systeme GmbH, Leutkirch im Allgäu, Germany) at increasing temperatures of 90 ℃ (2:30 min), 150 ℃ (5 min) and 210 ℃ (22:30 min). Phosphate-free filter paper (MN 280 ¼, Macherey-Nagel, Düren, Germany) was used to filter the extracts. The filtered extracts were used to measure the elements by ICP-OES (iCAP 7400 Series ICP–OES, Thermo Fisher Scientific, Dreieich, Germany) at the following wavelengths (nm): 589.592 (radial) for Na, 766.490 (radial) for K, 317.933 (axial) for Ca, 285.213 (axial) for Mg, 260.569 (axial) for Mn, 238.204 (axial) for Fe, 308.215 (axial) for Al, 182.034 (axial) for S and 185.942 (axial) for P. The calibration was performed using element concentrations of 1 g l^-1^ (Einzelstandards, Bernd Kraft, Duisburg, Germany) and an internal mixed standard.

### 2.3. DNA extraction and Illumina sequencing

Frozen soil samples (-20 ℃) were milled in liquid nitrogen using a ball mill (MN400, Retsch GmbH). About 250 mg of frozen, milled soil was used for the extraction of DNA with the DNeasy® PowerSoil® Pro kit (Qiagen, Hilden, Germany) and purified with the DNeasy® PowerClean® kit (Qiagen) according to the manufactureŕs recommendations. The concentration of extracted DNA was quantified using the NanoDrop 2000 spectrophotometer (PEQLAB Biotechnologie GmbH, Erlangen, Germany). The internal transcribed spacer (ITS) region was amplified by polymerase chain reaction (PCR) using the forward primer ITS3KYO2 (Toju et al., 2012) and reverse primer ITS4 (White et al., 1990). The PCR reactions, purification, pooling of triplicate technical replicates, and quantification of amplicons were conducted as described previously (Clausing et al. 2020). The amplicons were sequenced with MiSeq Reagent Kit v3 (Illumina Inc., San Diego, USA) using the MiSeq platform at the Göttingen Genomics Laboratory (G2L).

### 2.4. Bioinformatics data processing

Raw paired-end sequences were subjected to quality filtering using fastp V0.20.0 (Chen et al., 2018) and merged using PEAR v0.9.11 (Zhang et al., 2014). Additional forward and reverse primer clipping was performed with cutadapt v2.5 (Martin, 2011). VSEARCH v2.14.1 (Rognes et al., 2016) was used for the removal of reads with length <140 bp, length sorting, dereplication, denoising and exclusion of chimeric DNA sequences de novo and reference-based using the UNITE dataset (UNITE+INSDC v8.2, https://plutof.ut.ee/#/doi/10.15156/BIO/786370) accessible at https://unite.ut.ee/repository.php (Nilsson et al., 2015). To generate the operational taxonomic units (OTUs), the amplicon sequence variants (ASVs) were clustered at 97 % identity threshold using VSEARCH (--sortbysize and --cluster_size) and mapped to OTU sequences with VSEARCH (--usearch_global, -id 0.97). The resulting OTUs were taxonomically classified using BLAST 2.9.0+ against UNITE+INSDC v8.2 database (Kõljalg et al., 2013). All unidentified OTUs and OTUs with no blast hits were aligned against the nt database of the GenBank (Geer et al., 2010) and only OTUs with fungal classification were kept. The count number of fungal sequences per soil sample ranged from 13,668 to 108,070. The OTUs were assigned to fungal trophic modes (symbiotrophs, saprotrophs and pathotrophs) based on the FUNGuild database (Nguyen et al., 2016). The resulting OTU table was rarefied to 13,668 reads per sample (i.e. minimum number of reads present in one of our 120 samples) using the rrarefy() function in R vegan package v2.5.7 (Oksanen et al., 2020). For further analyses, the three replicate samples per plot were pooled yielding an OTU table with 40 samples. The data matrix is available in PANGEA (Likulunga et al., 2021).

The raw sequences were deposited in the National Center for Biotechnology Information (NCBI) Sequence Read Archive (SRA) (Leinonen et al., 2011) with Bioproject accession number PRJNA704813.

### 2.5. Statistical analysis

The statistical analyses were performed with R software version 4.0.2 (R Core Team, 2020). Normal distribution and homogeneity of variances were tested using model residuals by conducting the Shapiro Wilk test. If the normality assumption was not met, data were subjected to square root or log transformation. If data sets were not normal-distributed after transformation, non-parametric tests (Kruskal-Wallis test) were used. Under normality assumptions, analysis of variance (ANOVA) was performed followed by a post hoc Tukey HSD test for comparison of means (package: “multcomp”). Differences of the means were considered to be significant at p ≤ 0.05.

Non-metric multidimensional scaling (NMDS) with Bray Curtis as metric distance was used to visualize the dissimilarities of soil fungal communities. Analysis of similarity (ANOSIM) (package: “vegan”) (Oksanen et al., 2020) with 9999 iterations was performed to determine significant differences of soil fungal communities among tree stand types. Pearson’s pairwise correlation matrix was used to determine collinearity among variables using the “Hmisc” package (Harrell et al., 2020) and variables with high correlation coefficients (r > 0.8) were not used for analysis with exception of N, P and CN ratio. We used the “envfit function” in the “vegan” package (Oksanen et al., 2020) to correlate the soil properties (moisture, concentration of total phosphorus, total calcium, and nitrogen) and proportion of conifers to soil fungal communities.

We used general linear and linear mixed effect models with site as random factor and applied the post hoc Tukey HSD test to test for differences at p ≤ 0.05 of species richness and diversity of soil fungal communities and fungal modes (symbiotrophs (SYM), saprotrophs (SAP) and pathotrophs (PAT)) among the tree stands. Effect size for a distinct stand type relative to beech was determined for the SYM fungi was determined as follows:

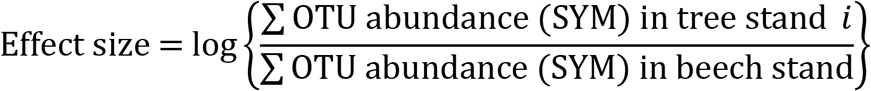

where tree stand *i* refers to a specific stand type (such as Douglas-fir, spruce, mixture of beech with Douglas-fir or mixture of beech with spruce). The effect size of saprotrophic fungi was determined accordingly. The effect sizes were analyzed for differences among tree stands using linear mixed effect models with site as random factor and applying the post hoc Tukey HSD test. We used One-sample Wilcoxon signed-rank test (tested against median = 0) to test for differences of effect size from zero in each tree stand.

We used Linear Discriminant Analysis (LDA) analysis to test for differences among sites by soil properties and forest types using the “MASS” R package (Venables and Ripley, 2002). Generalized Adaptive Models for Location, Scale and Shape with a zero-inflated beta family (GAMLSS-BEZI) from the “metamicrobiomeR” package (Ho et al., 2019) were used to test for differences of the relative abundances of a distinct group of fungi between tree stands. GAMLSS is a type of regression models in which the response variable can take any type of distribution (Rigby and Stasinopoulos, 2005), and with zero-inflated beta (BEZI) family, the model performs efficiently in analyzing relative abundance data, providing the meta-analysis output is based on log of odds ratio (Ho et al., 2019). Therefore, GAMLSS-BEZI is suitable for analysis of relative abundance data.

## 3. Results

### 3.1. Fungal diversity in soil of different stand types and sites

We sequenced 120 soil samples and obtained a total number of 4,941 million sequence reads, which clustered into 2,198 OTUs (potential species). Fungal OTU richness showed significant differences among the stand types (p < 0.001, Chi² = 85.39), with Douglas-fir stands exhibiting the highest (609 ± 42) and beech stands the lowest OTU richness (514 ± 26) (Fig. 1a). Mixed stands of beech and conifers had higher species richness than pure beech stands and the fungal OTU richness of spruce stands was intermediate compared to beech and Douglas-fir (Fig. 1a).

**Fig 1.**
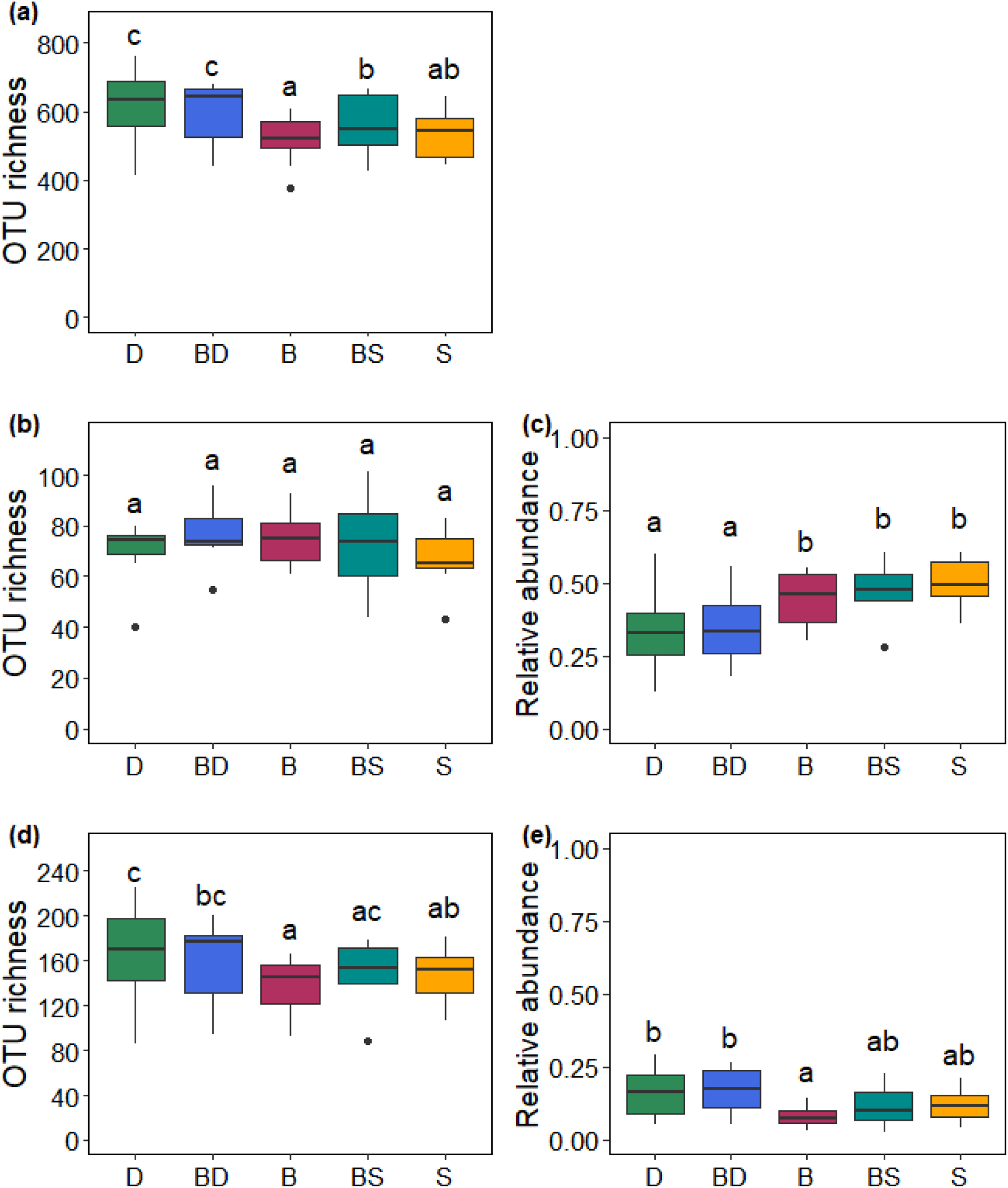
Fungal species richness and relative abundance in all (a), symbiotrophic (b and c) and saprotrophic (d and e) fungi in soil of beech, conifer and mixed stands. Douglas-fir (D), beech (B), spruce (S), mixture of beech with Douglas-fir (BD) and mixture of beech with spruce (BS). Lower case letters indicate significant differences at p ≤ 0.05 (n = 8) among the tree stands using linear mixed effect and Kruskal-Wallis test.

In order to investigate whether stand type affected fungal richness in distinct trophic groups, we analyzed OTU richness of symbiotrophs and saprotrophs. Symbiotrophic fungi, a group which was formed here mainly by ectomycorrhizal fungi (83%, read abundance table: Likulunga et al., 2021), did not show any differences in species richness (74 ± 3) across the five stand types studied here (Fig. 1b). However, in contrast to species richness, the relative abundances (based on sequence reads) of the symbiotrophic fungi varied among forest types and were lowest in the Douglas-fir and beech-Douglas-fir stands, intermediate in beech and beech-spruce mixture and highest in spruce stands (Fig. 1c).

The richness of saprotrophic fungi varied significantly among the stand types, with pure Douglas-fir stands having the highest (166 ± 15) and beech stands the lowest OTU richness of saprotrophic fungi (137 ± 9) (Fig. 1d). The relative sequence abundance of the saprotrophic fungi was lowest in beech and highest in Douglas-fir stands (Fig. 1e). Overall, the relative sequence abundance of the saprotrophic fungi was much lower than that of the symbiotrophic fungi (p < 0.001, Chi² = 26.14) (Fig. 1c, e; Supplement Fig. S1).

Fungal OTU richness was not only affected by stand type but also by site (p < 0.001, Deviance = 375.53) and location of the study sites (p < 0.001, Deviance = 12.06).

Similarly, Shannon diversity showed differences among the sites, an effect that was due to significantly lower Shannon diversity of symbiotrophic fungi in beech-spruce mixture than in beech or Douglas-fir stands (Table 1). The Shannon diversity of the saprotrophic fungi was unaffected by stand type and site (Table 1).

**Table 1.**
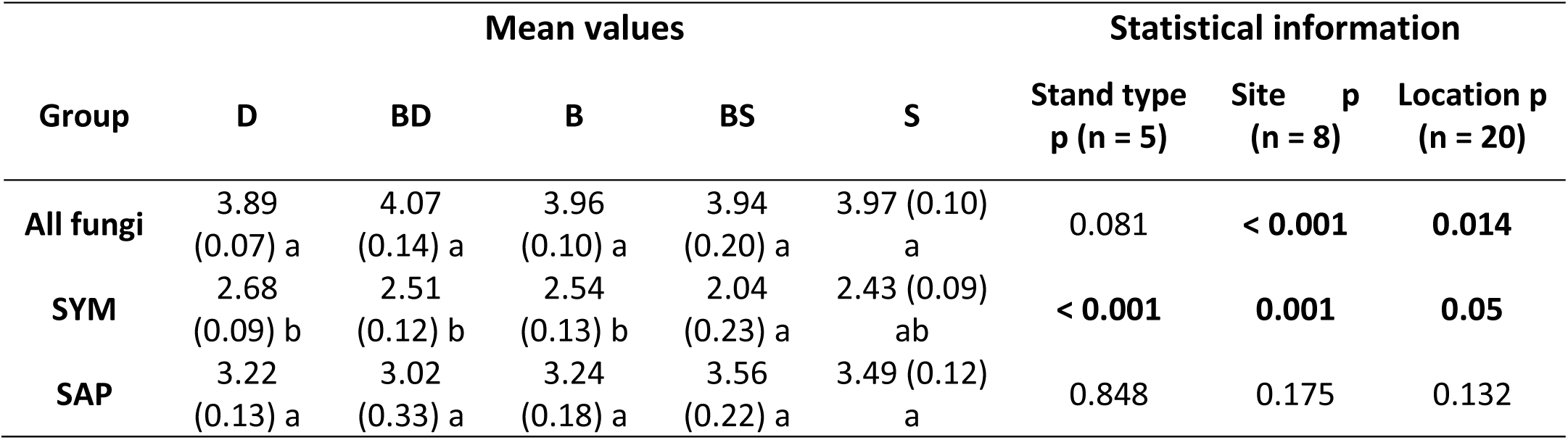
Shannon diversity of soil fungal communities for all, symbiotrophic (SYM) and saprotrophic (SAP) fungi in different stand types, study sites and locations of the study sites. Data are indicated as means (± SE) for stands of Douglas-fir (D), beech (B), spruce (S), mixture of beech with Douglas-fir (BD) and mixture of beech with spruce (BS). Means among tree stand types, sites and locations were compared by linear models and post hoc Tukey HSD test. Different letters in rows indicate significant differences at p ≤ 0.05.

We also evaluated the effect sizes of the changes in fungal abundances relative to pure beech stands (Fig. 2). A significant, negative effect of Douglas-fir and beech-Douglas-fir stands on symbiotrophic fungal abundances compared to the native spruce and beech-spruce stands (p < 0.001, Chi² = 24.40) was recorded, (Fig. 2a). In spruce or beech-spruce stands, no significant effects were observed (Fig. 2a). The effect sizes of saprotrophic fungi in conifer or mixed beech-conifer stands did not differ significantly among each other (p = 0.145, Chi² = 5.40) (Fig. 2b). However, we observed significantly higher effect sizes of saprotrophic fungi in Douglas-fir (p = 0.030), spruce (p = 0.042) and beech-Douglas-fir (p = 0.014) stands compared with pure beech, which resulted in positive effects (Fig. 2b).

**Fig 2.**
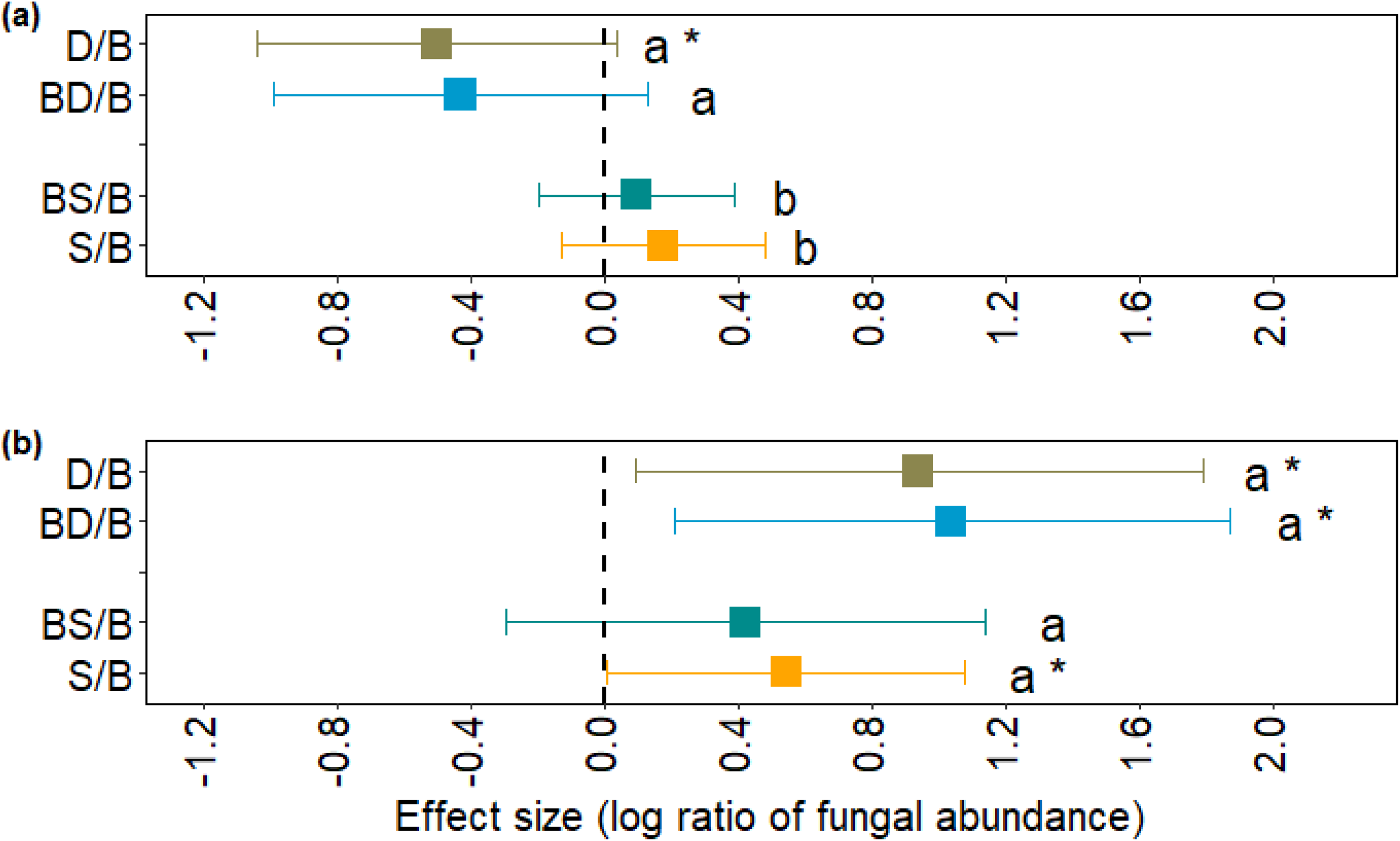
Effect size for symbiotrophic (a) and saprotrophic (b) fungal abundance in conifer and mixed tree stands relative to pure beech stands (D/B: Douglas-fir and beech; BD/B: Mixture of beech with Douglas-fir and beech; BS/B: Mixture of beech with spruce and beech; S/B: spruce with beech). Effect size refers to the log ratio of fungal abundance in conifers and beech-conifer mixtures to beech stands. Lower case letters indicate significant differences among tree stand at p ≤ 0.05 (ANOVA for linear mixed effect model with site as random factor (n = 8)). Significant differences (p < 0.05) of effect sizes from zero are indicated by * (One-sample Wilcoxon signed-rank test).

### 3.2 Soil chemistry in different stand types and sites

Soil properties (relative soil moisture, pH, C, N, CN ratio, Ca, Mg, K, Na, Mn, Fe, S, P and Al) varied significantly among the different stand types and sites (Table 2). We tested whether these properties could be used to discriminate between stand types and sites. Linear Discriminant Analysis (LDA) of all measured soil properties did not reveal a clear differentiation among different forest types (Fig. 3a, confusion matrix in Supplement Table S1). The separation of sites was more distinct (Fig. 3b) but the confusion matrix showed that only four out of eight sites were correctly assigned (Supplement Table S1). The best separation was obtained when the sites were grouped according to location in the southern (Harz, Dassel, Winnefeld, Nienover) and northern region (Nienburg, Unterlüss, Göhrde I, Göhrde II, Fig. 3c) of our study area. This separation also reflected different soil types with clay-silt soils in the south and sandy soils in the north (Foltran et al., 2020).

**Fig 3.**
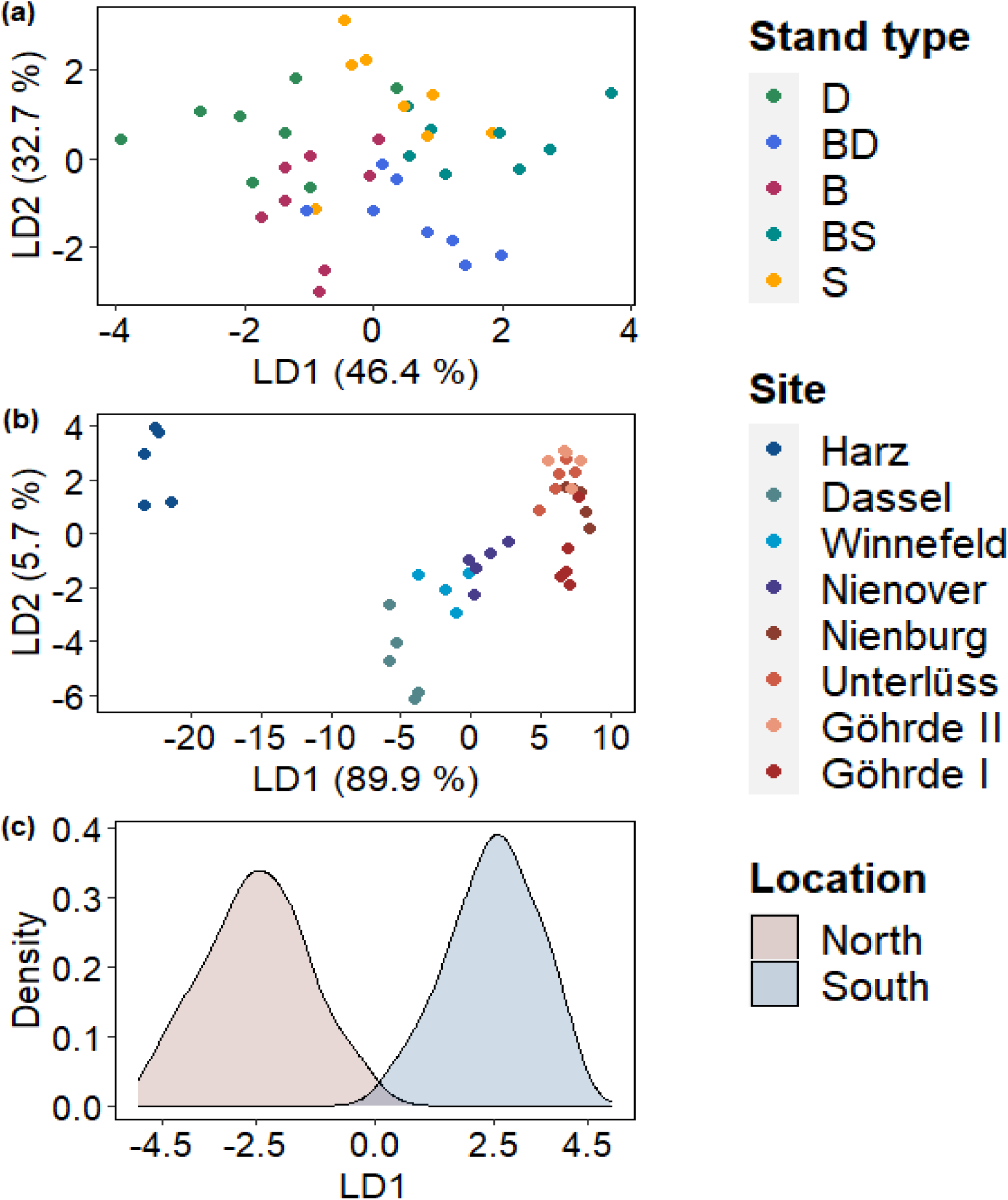
Linear discriminant analysis (LDA) according to stand types (a), study sites (b) and location of the study sites (c). The analysis was conducted on measured soil properties (relative soil moisture, pH, C, N, CN ratio, Ca, Mg, K, Na, Mn, Fe, S, P and Al). The corresponding confusion matrix is provided as Supplement Table S3.

**Table 2.** Soil properties in different tree stand types, study sites and locations of the study sites. Data are indicated as means (± SE) for stands of Douglas-fir (D), beech (B), spruce (S), mixture of beech with Douglas-fir (BD) and mixture of beech with spruce (BS). Means among tree stand types, sites and locations were compared by linear models and post hoc Tukey HSD test. Different letters in rows indicate significant differences at p ≤ 0.05.

| Variable | Mean values in stand types |  |  |  |  | Statistical information |  |  |
| --- | --- | --- | --- | --- | --- | --- | --- | --- |
|  | D | BD | B | BS | S | Stand type<br>p (n = 5) | Site<br>p (n = 8) | Location<br>p (n = 20) |
| **pH** | 3.47<br>(0.09) b | 3.41<br>(0.08) b | 3.40<br>(0.06) b | 3.18<br>(0.05) a | 3.30<br>(0.08) ab | **0.001** | **< 0.001** | **< 0.001** |
| **MC** | 38.27<br>(2.56) a | 45.26<br>(3.30) ab | 42.07<br>(2.74) a | 54.71<br>(4.40) c | 50.65<br>(2.69) bc | **< 0.001** | **< 0.001** | **< 0.001** |
| **CN** | 23.89<br>(0.62) b | 22.01<br>(1.00) a | 20.53<br>(0.52) a | 23.43<br>(0.68) b | 23.54<br>(0.76) b | **< 0.001** | **< 0.001** | **< 0.001** |
| **C** | 63.08<br>(6.99) ab | 75.05<br>(9.23) bc | 52.14<br>(4.17) a | 81.10<br>(6.10) c | 75.45<br>(4.39) bc | **< 0.001** | **< 0.001** | 0.981 |
| **N** | 2.63<br>(0.26) a | 3.34<br>(0.33) ab | 2.57<br>(0.19) a | 3.58<br>(0.31) b | 3.25<br>(0.22) b | **< 0.001** | **< 0.001** | **< 0.001** |
| **Na** | 0.18<br>(0.03) ab | 0.23<br>(0.05) b | 0.16<br>(0.02) ab | 0.13<br>(0.01) a | 0.20<br>(0.03) b | **0.008** | **< 0.001** | **< 0.001** |
| **K** | 0.77<br>(0.16) b | 0.84<br>(0.16) b | 0.77<br>(0.17) ab | 0.45<br>(0.07) a | 0.64<br>(0.12) (ab) | **0.001** | **< 0.001** | **< 0.001** |
| **Ca** | 0.89<br>(0.06) ab | 1.01<br>(0.12) b | 0.79<br>(1.10) a | 0.76<br>(0.05) ab | 0.91<br>(0.06) ab | **0.022** | **< 0.001** | **< 0.001** |
| **Mg** | 1.28<br>(0.25) bc | 1.48<br>(0.26) c | 1.31<br>(0.31) ab | 0.78<br>(0.12) a | 1.14<br>(0.18) bc | **< 0.001** | **< 0.001** | **< 0.001** |
| **Mn** | 0.14<br>(0.02) a | 0.26<br>(0.05) a | 0.20<br>(0.04) a | 0.16<br>(0.04) a | 0.15<br>(0.03) a | 0.071 | **< 0.001** | **< 0.001** |
| **Fe** | 9.88<br>(1.74) a | 10.02<br>(1.17) a | 9.07<br>(1.78) a | 6.51<br>(0.68) a | 8.54<br>(1.38) a | 0.240 | **< 0.001** | **< 0.001** |
| **S** | 0.34<br>(0.03) ab | 0.36<br>(0.04) ab | 0.28<br>(0.02) a | 0.35<br>(0.02) ab | 0.39<br>(0.03) b | **0.004** | **< 0.001** | 0.734 |
| **P** | 0.27<br>(0.03) a | 0.31<br>(0.03) ab | 0.28<br>(0.03) ab | 0.28<br>(0.03) a | 0.34<br>(0.04) b | 0.010 | < 0.001 | < 0.001 |
| **AI** | 15.90<br>(3.25) b | 15.29<br>(3.01) ab | 14.47<br>(3.26) ab | 9.27<br>(1.23) a | 13.44<br>(2.53) ab | 0.023 | < 0.001 | < 0.001 |

### 3.3. Soil fungal communities cluster according to stand type and soil chemistry

To determine whether the composition of soil fungi varied with stand type and soil properties, we visualized the dissimilarities of fungal assemblages with an NMDS (Fig. 4 a). The first ordination axis separated the fungal communities according northern and southern locations (ANOSIM for location: R = 0.346, p <0.0001). The separation along the first axis was accompanied by changes in soil chemistry (pH, N, C/N ratio, P, Ca and soil moisture, Fig. 4b, Supplement Table S2).

**Fig. 4.**
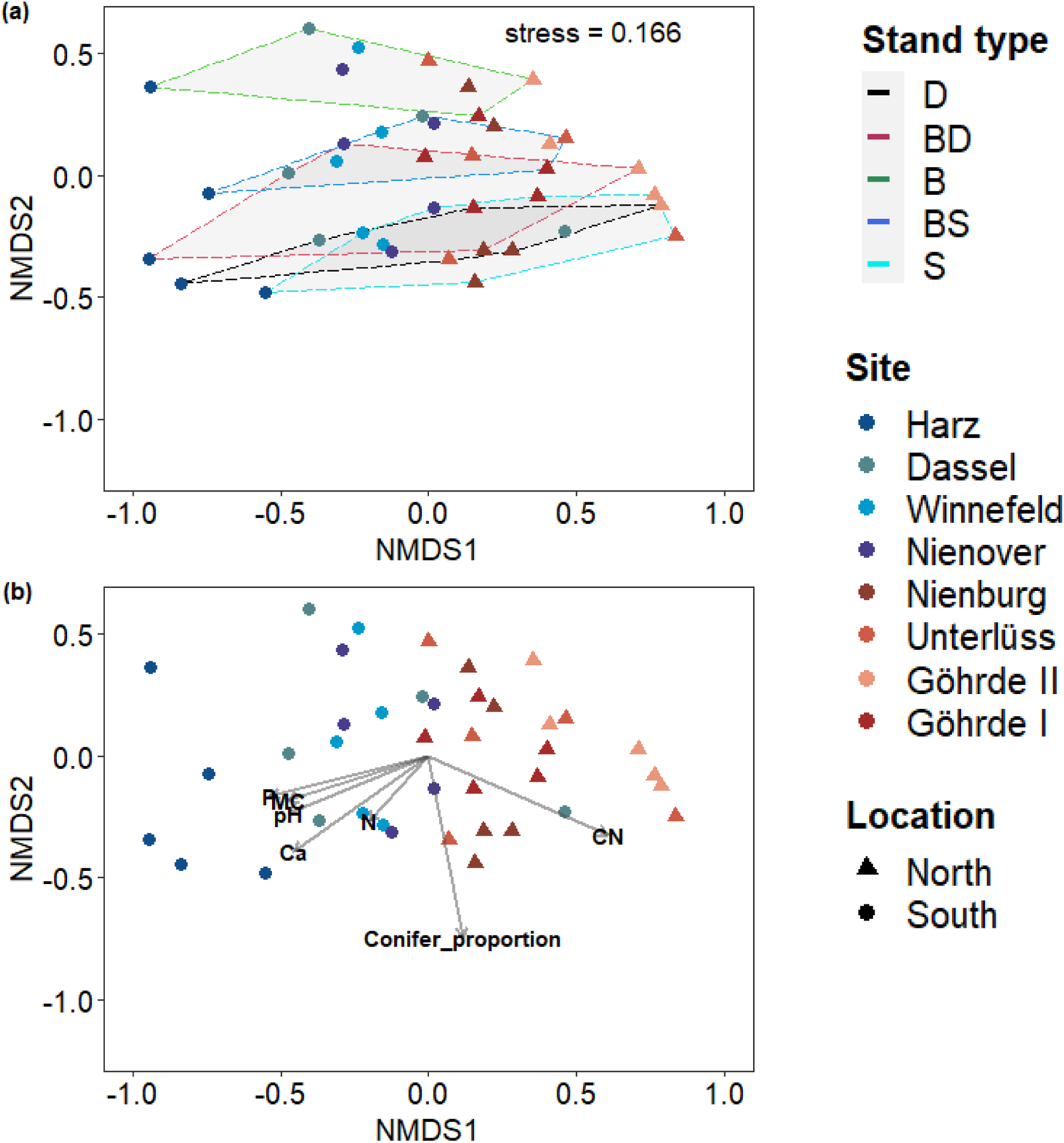
Non-metric multidimensional scaling (NMDS) of soil fungal OTU communities based on Bray-Curtis dissimilarity measure (a) and main explanatory variables (b). The tree stands in (a) are indicated by hulls: beech (B), spruce (S), Douglas-fir (D), mixture of beech with spruce (BS) and mixture of beech with Douglas-fir (BD). Significant vectors (b) indicate: soil moisture (MC), concentration of calcium (Ca), concentration of phosphorous (P), soil pH, concentration of nitrogen (N), CN ratio and proportion of conifers (Conifer_proportion). Data points indicate sum of OTUs (n =3) and means of soil properties (n = 3) per site and forest types).

The second axis separated the fungal community composition according to stand type (Fig. 4a, ANOSIM for the stand types: R = 0.291 and p < 0.0001) and was related to the proportion of conifers (Fig. 4b, Supplement Table S2). The composition of the fungal communities in soil of beech stands clustered separately from those in mixed and those in conifers stands (Fig. 4a). The fungal community compositions of spruce and Douglas-fir stands showed strong overlap and those of the mixture with beech were intermediate to the pure stands (Fig. 4a)

### 3.3. Tree stand type and site show shifts in fungal orders

Since we found that the separation of the fungal species (OTUs) was mainly driven by stand type and location (north, south), the question arose whether this pattern was caused by loss or appearance of distinct phylogenetic groups of the fungi. To address this question, OTUs were grouped at the level of fungal orders. We used all fungal orders (n = 16) with relative abundances above 1% of the sequences for our analysis (Supplement Table S3, Supplement Fig. S2). These orders encompassed together more than 80% of the sequences (Supplement Fig. S2, Supplement Table S4).

Russulales, an ectomycorrhizal fungal order (Rinaldi et al., 2008; Tedersoo et al., 2010), was enriched in beech stands in the south but not in the north locations compared to Douglas-fir (Fig. 5a). Therefore, no difference for the Russulales between beech and Douglas-fir forests was observed when the south and north locations were combined (Fig. 5a). None of the other fungal orders showed a significant shift between Douglas-fir and beech in the north or south locations (Fig. 5a). However, the pooled data for both locations revealed that Tremellales (mainly mycoparasitic yeasts, Sterkenburg et al., 2015), Hymenochaetales (containing important forest pathogens and wood degrading fungi, Tedersoo et al., 2014) and Helotiales (saprotrophic and ectomycorrhizal fungi, Cannon and Kirk, 2007) increased in Douglas-fir stands, while Boletales (mainly ectomycorrhizal fungi, Cannon and Kirk, 2007) and Agaricales (ectomycorrhizal and saprotrophic fungi, Kirk et al., 2008) were enhanced in beech stands (Fig. 5a).

**Fig 5.**
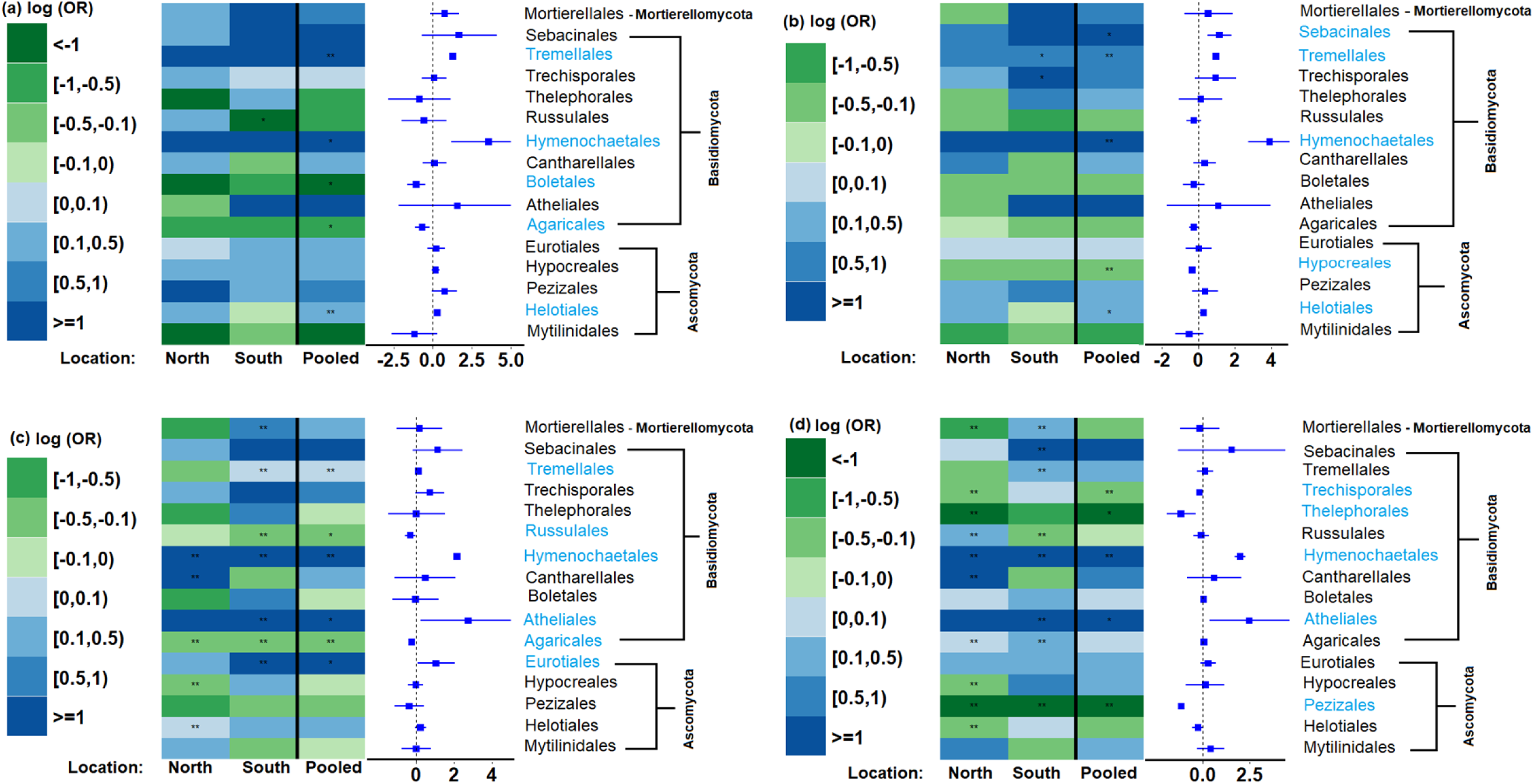
Changes in the relative abundance of fungal orders in conifer and mixed stands compared with beech:(a) Douglas- fir/ beech, (b) mixture of beech with Douglas-fir/ beech, (c) spruce/beech, (d) mixture of beech with spruce/beech. Data are shown as log ratio of odds (log(OR)). Green cells (negative values) indicate enrichment of a fungal order in beech forests and blue (positive values) indicate enrichment in the conifer forest or mixture. Significant differences at p ≤ 0.05 (GAMLSS model with BEZI family, n = 4 for the fungal orders in stands in the north and south location, n = 8 for the pooled data) are indicated with stars (* p < 0.05, ** p < 0.01). Significant orders are highlighted in blue.

In the mixed beech-Douglas-fir stands, Tremellales, Hymenochaetales and Helotiales were also enriched compared with beech stands (Fig. 5b). Furthermore, Sebacinales (many ectomycorrhizal and saprotrophic species, Cannon and Kirk, 2007) increased in the mixed beech-Douglas-fir stands, while Hypocreales (saprotrophic and pathogenic species, Sterkenburg et al., 2015) were enriched in beech stands (Fig. 5b). In spruce and spruce-beech mixtures Tremellales (under most conditions) and Hymenochaetales were enriched compared to beech, suggesting that these orders prefer conifer forests (Fig. 5c,d). In the south locations, Russulales were enriched in beech forests compared to spruce and Atheliales (saprotrophic and mycorrhizal species, Jülich, 1918; Kirk et al., 2008) in spruce and mixed forests compared to beech (Fig. 5c,d). Pezizales (many saprotrophic and ectomycorrhizal species, Cannon and Kirk, 2007) were strongly enriched across all sites in beech forests compared with spruce-beech mixtures (Fig. 5d). Overall, a higher number of fungal orders showed significant variation in spruce or beech-spruce mixture compared with beech than Douglas-fir and its mixture (Fig. 5).

We also compared the changes in fungal relative abundances between spruce and Douglas-fir stands and between spruce-beech and Douglas-fir beech stands (Supplements Fig. S3a,b). The spruce-Douglas-fir comparison showed a number of significantly affected fungal orders similar to that found in the beech comparisons (i.e. 5 or 6 out of 16 orders) (Supplement Fig. S3a). The comparison of the mixtures showed the highest number of significant variations (11 out 16) in our study (Supplement Fig. 3b).

### 4. Discussion

### 4.1. Fungal taxonomic community composition is strongly driven by site conditions and forest type

Our results show that Douglas-fir, spruce, beech and the mixtures of beech with these conifer species shaped the fungal communities in soil. These results are in agreement with other studies demonstrating that tree species or community effects influence the structure of soil fungal assemblages (Prescott and Grayston 2013; Urbanova et al., 2015; Nacke et al., 2016; Bahmann et al. 2018; Prada-Salcedo et al., 2020). Our results further support that main environmental drivers of fungal community composition are soil C/N, pH and soil moisture (Wubet et al., 2012; Kivlin et al., 2014; Sterkenburg et al., 2015; Glassman et al., 2017; Schröter et al., 2019; Vetrovský et al., 2019; Clausing et al., 2020; Nguyen et al., 2020). These results show that the overall site effects found in our study concur with those reported previously and explain the separation of the fungal communities between the northern and the southern region, which are characterized by strong differences in soil properties, soil resources and climate (Ammer et al., 2020; Foltran et al., 2020; this study).

A major goal was to understand better the influence of an introduced conifer compared with indigenous tree species on soil fungi. In contrast to our first hypothesis, we did not observe a reduction but an increase in fungal species richness under Douglas-fir. Since the Douglas-fir stands in our study had an age of several decades (Ammer et al., 2020) and Douglas-fir has been introduced to Germany about 150 years ago (Essl, 2005), we speculate that adapted fungal species might have evolved the ability to form novel associations. Another explanation is that ecologically compatible fungal species occur in Douglas-fir biomes because of the wide geographic range of fungal species (Tedersoo et al., 2014). Our results support the latter assumption as the fungal community structures in Douglas-fir and spruce forest soils showed a strong overlap. This notion does not exclude that evolutionary processes are taking place but due to the similarity of the fungal composition of Douglas-fir and spruce found here, putative new or adapted fungal species or strains were subordinate and not traceable with marker gene-based approaches.

Here, we report a clear separation of fungal composition between beech and conifers while beech-mixed stands exhibited intermediate soil fungal assemblages. Previous studies also found differences in soil fungal communities between beech and spruce forests (Sterkenburg et al., 2015; Bahnmann et al., 2018; Nacke et al., 2016; Asplund et al., 2019). The shifts observed in soil fungal communities among different forests types have been attributed to indirect effects of dominant tree species through modification of soil resources (Augusto et al., 2002; Augusto et al., 2015). Coniferous ecosystems show lower foliage nutrients (e.g. Ca, K, Mg, P and N) than ecosystems dominated by broadleaf species (Augusto et al., 2002; Augusto et al., 2015), whereby soil nutrients can be influenced through litter production (Stoutjesdijk and Barkman, 1992), consequently leading to tree-driven soil fungal communities (Prescott and Grayston 2013; Urbanová et al., 2015). For example, Asplund et al. (2019) found that the fungal communities in beech and spruce forests were separated by pH and the C/N ratio. Similarly, Nacke et al. (2016) reported that pH and organic carbon were major drivers for the separation of beech and spruce fungal communities. However, Bahnmann et al. (2018) showed that pH and C/N affected only the ectomycorrhizal fungi in beech and spruce forests within a distinct geographic area, while soil moisture was a main factor at larger scales. In our study, the abiotic factors (pH, moisture) separated the fungal communities in two regions, each containing all forest types. This result highlights the importance of the geographic scale of the studies. We found that the separation according to forest types was mainly influenced by the fraction of conifers. Spruce and Douglas-fir forests contain higher and beech-conifer mixed stands intermediate organic C and N stocks compared to beech stands (Cremer et al., 2016). Differences in litter resources might explain the distinct assemblages observed between beech, conifers and their mixtures. However, there is now increasing evidence that root exudates also play profound roles in shaping belowground microbial communities because of differences in quantity and quality among tree species (Brimecombe et al., 2000; Bertin et al., 2003; Jones et al., 2004; Haichar et al., 2014; Kardol and De Long, 2018). Experimental studies are required to clarify these points.

### 4.2 Douglas-fir affects the functional composition of fungal communities

We discovered functional differences in the fungal assemblages among distinct forest types with the highest species richness of saprotrophic fungi in Douglas-fir soils. Furthermore, the relative abundance of saprotrophic fungi was higher in Douglas-fir and beech-Douglas-fir mixture than in beech stands, which might be attributable to litter chemistry, known to shape the identity and composition of saprotrophs (Kubartová et al., 2009; Treseder et al., 2014; Urbanová et al., 2015; Foudyl-Bey et al., 2016; Bahnmann et al., 2018). Douglas-fir litter contains high cellulose and low lignin contents (Kubartová et al., 2009), consequently favoring the decomposer communities. This may explain the higher proportions of saprotrophs observed in Douglas-fir than beech stands. Further, Awad et al. (2019) found a strong influence of conifers on saprotrophic fungal biomass, also linked to soil resources such as N and C. In spruce forests, the relative abundance of saprotrophic fungi was intermediate between beech and Douglas-fir, while the relative abundance of symbiotrophic fungi was higher in beech and spruce than in Douglas-fir forests. These shifts affected the balance between saprotrophic and symbiotrophic fungi in a complex manner, resulting in positive effect sizes for saprotrophic fungi in the conifer forests but a specific negative effect size for symbiotrophic fungi in Douglas-fir soil. These patterns might reflect inter-guild interactions in which saprotrophs and symbiotrophs compete for soil resources, hence inhibiting each other and slowing down decomposition (Gadgil and Gadgil, 1975; Leake et al., 2002; Fernandez & Kennedy, 2016). Since forest types clearly affected the composition with increases in the fungal decomposer community favored especially by Douglas-fir, we conclude that tree identity effects shape the functional composition of fungal assemblages.

### 4.3 Forest type and site conditions have different impact on genetically related fungal taxa

We gained insights into the responsiveness of fungal taxa (grouped at the rank of orders) to forest types (relative to beech) and regional factors. Several saprotrophic fungal orders showed variability between north and south locations, indicating susceptibility to microclimate and soil resource. However, one order was specifically enriched in stands with Douglas-fir (Helotiales), and one (Eurotiales) in pure spruce stands. Furthermore, there were also common positive effects for wood degrading fungi and yeasts (Hymenochaetales, Tremellales, Tedersoo et al., 2014; Sterkenburg et al., 2015) across all site conditions and conifers. This result shows that both spruce and Douglas-fir foster similar saprotrophic fungal groups, thus, contributing to the observed increase of saprotrophic potential. Genome sequencing of members of the Hymenochaetales demonstrated a high number of genes for carbohydrate-active enzymes in the analyzed species (Kohler et al. 2015). Thus, our results underpin that genomic information can enlighten ecological processes.

In temperate beech stands, members of the Russulales are usually highly abundant (Tedersoo et al., 2014; Buée et al. 2005; Lang et al. 2011; Pena et al. 2017). Russulales are a ubiquitous fungal group, in which ectomycorrhizal fungi dominate (Looney et al. 2018). Russulales were the most abundant fungal group in our study. They appear to be specialized for the acquisition of ammonium (Nygren et al. 2008). Since they descent from white-rot fungi, Russulales may be able to attack lignin (Looney et al. 2018) but may not be capable to accessing C from cellulose and other C-rich biopolymers (Wolfe et al., 2012). In agreement with other studies (Uroz et al., 2016; Asplund et al., 2019), we found that members of the Russulaceae were enriched in beech compared to spruce forests. However, this result was only obtained in pure forests, whereas in mixed or Douglas-fir forests, positive effects were only observed in the south, where soil N contents were higher than in the north. Thus, variation of the Russulales appears to be driven by resource availability. Similarly, Boletales, which contain ectomycorrhizal fungi with oxidative enzymes able to degrade lignin (Op De Beeck et al. 2018), were enriched in beech compared with Douglas-fir but were variable in mixed and spruce forests. Atheliales was the only ectomycorrhizal fungal order that showed a positive effect in a conifer forest.

Overall, the commonalities of Douglas-fir and spruce were confined to saprotrophic taxa, while the influence on other orders was context-dependent and affected saprotrophic and ectomycorrhizal guilds likewise. The importance of substrate quality for fungal populations has been highlighted (Allison et al., 2007; Bossuyt et al., 2001; Christensen, 1989; Frey et al., 2004; Schröter et al. 2019; Nguyen et al. 2020). The phylogenetic positions of the affected fungal orders and genomic information from reference species suggest that mechanistic understanding of fungal assemblies comes into reach.

### 5. Conclusion

Our study demonstrates that Douglas-fir, a non-native tree species in Europe, is able to integrate native soil fungal networks similar to native spruce but different from beech. Mixing of beech with either Douglas-fir or spruce formed intermediate soil fungal composition but in contrast to our initial hypothesis, mixtures did not result in enhanced fungal species richness in soil. On top of abiotic soil properties and climatic factors (pH, C/N ratio, Ca, P and soil moisture), which had a strong impact on fungal community composition, we uncovered distinct effects of forest types. Douglas-fir and beech- Douglas-fir stands demonstrated a shift in fungal functional guilds, exhibiting low proportions of symbiotrophic and high of saprotrophic fungi. The shifts in taxonomic and functional structures of soil fungi observed in the forest types and different site conditions can potentially affect vital ecosystem processes such as decomposition, C sequestration and nutrient cycling. Therefore, long-term research incorporating quantitative analyses of fungal traits, turnover of soil organic matter and nutrient retention is needed to unravel the ecological impact of beech enrichment with Douglas- fir and spruce.

## Acknowledgements

We thank R. Thoms and M. Reichel for assistance with sampling, T. Klein, G. Lehmann, and M. Smiatacz for processing samples, M. Franke-Klein for element analyses, T. Klein for DNA extraction and library preparation.

## Authors’ Contributions

A.P, L.E.L and C.A.R.P: Conceptualization, Methodology. Formal analysis: A.P and L.E.L. Investigation: L.E.L. Data curation: D.S and R.D. Writing-original draft preparation: L.E.L. Writing-review and editing: A.P, L.E.L, C.A.R.P, D.S and R.D. Supervision: A.P.

---

**Supplement Table S1.**
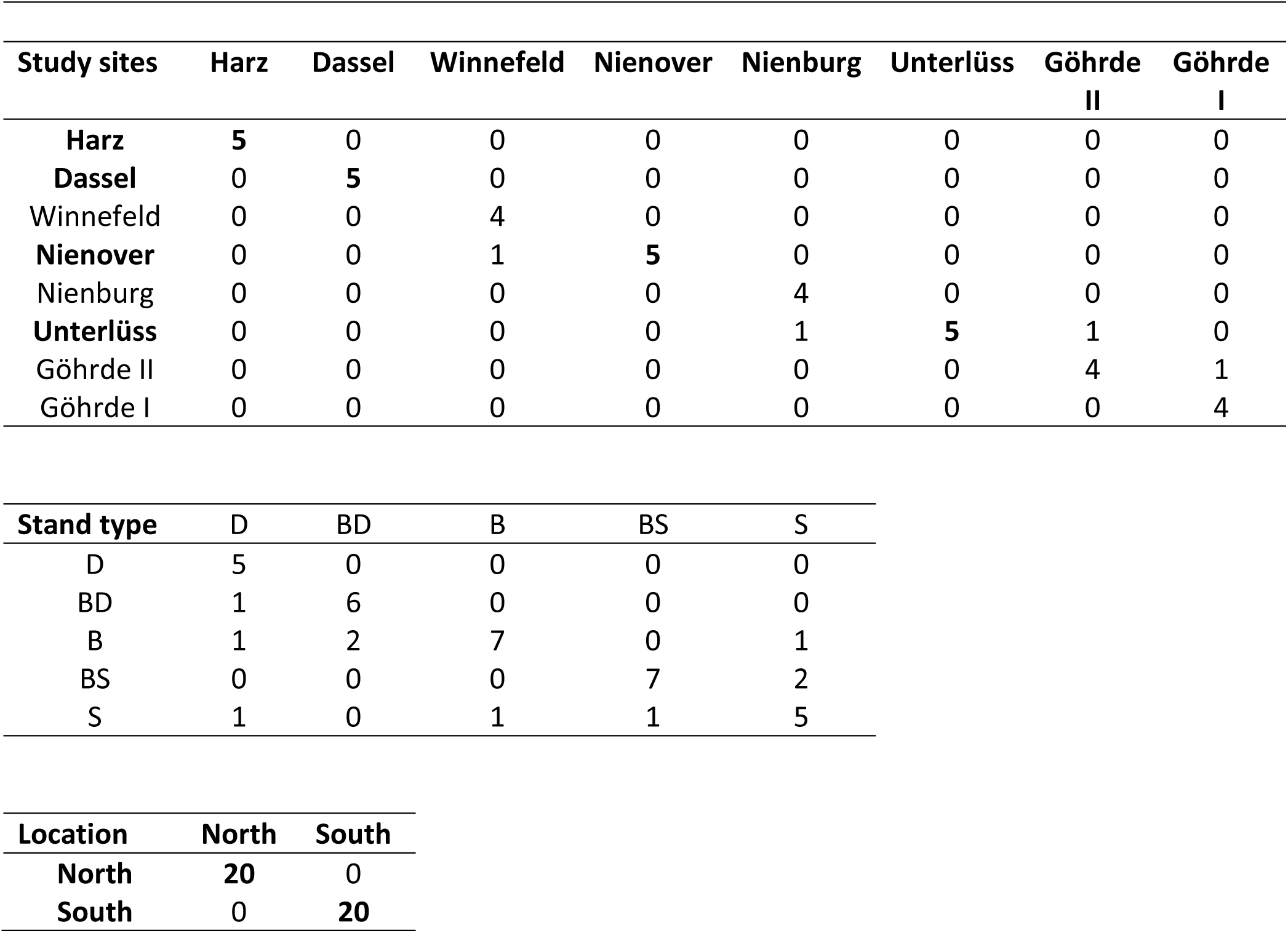
Confusion matrix based on linear discriminant analysis (LDA) of all measured soil properties. The plots with completely correct classification are indicated in bold.

**Supplement Table S2.** Correlation of explanatory variables to all, symbiotrophic and saprotrophic fungi based on envifit function. The explanatory variables soil properties (pH, MC: moisture, Ca: total calcium, P: total phosphorous, N: nitrogen and CN: CN ratio) and the proportion of conifers among the tree stands (Conifer_proportion) were included for analysis while variables C, Na, K, Mg, Fe, Mn and Al were excluded due to multicollinearity with exception of N, CN and P.

| Variable | All fungi |  | SYM |  | SAP |  |
| --- | --- | --- | --- | --- | --- | --- |
|  | R<sup>2</sup> | p | R<sup>2</sup> | p | R<sup>2</sup> | p |
| **pH** | 0.3931 | **0.002** | 0.3789 | **0.001** | 0.271 | **0.003** |
| **MC** | 0.3606 | **0.001** | 0.3944 | **0.001** | 0.4118 | **0.001** |
| **CN** | 0.6599 | **0.001** | 0.6524 | **0.001** | 0.6244 | **0.001** |
| **N** | 0.1568 | **0.046** | 0.1188 | 0.099 | 0.1978 | **0.015** |
| **Ca** | 0.5050 | **0.001** | 0.469 | **0.001** | 0.3604 | **0.003** |
| **P** | 0.4419 | **0.001** | 0.4556 | **0.001** | 0.5204 | **0.001** |
| **Conifer_proportion** | 0.7936 | **0.001** | 0.6701 | **0.001** | 0.7266 | **0.001** |

**Supplement Table S4.**
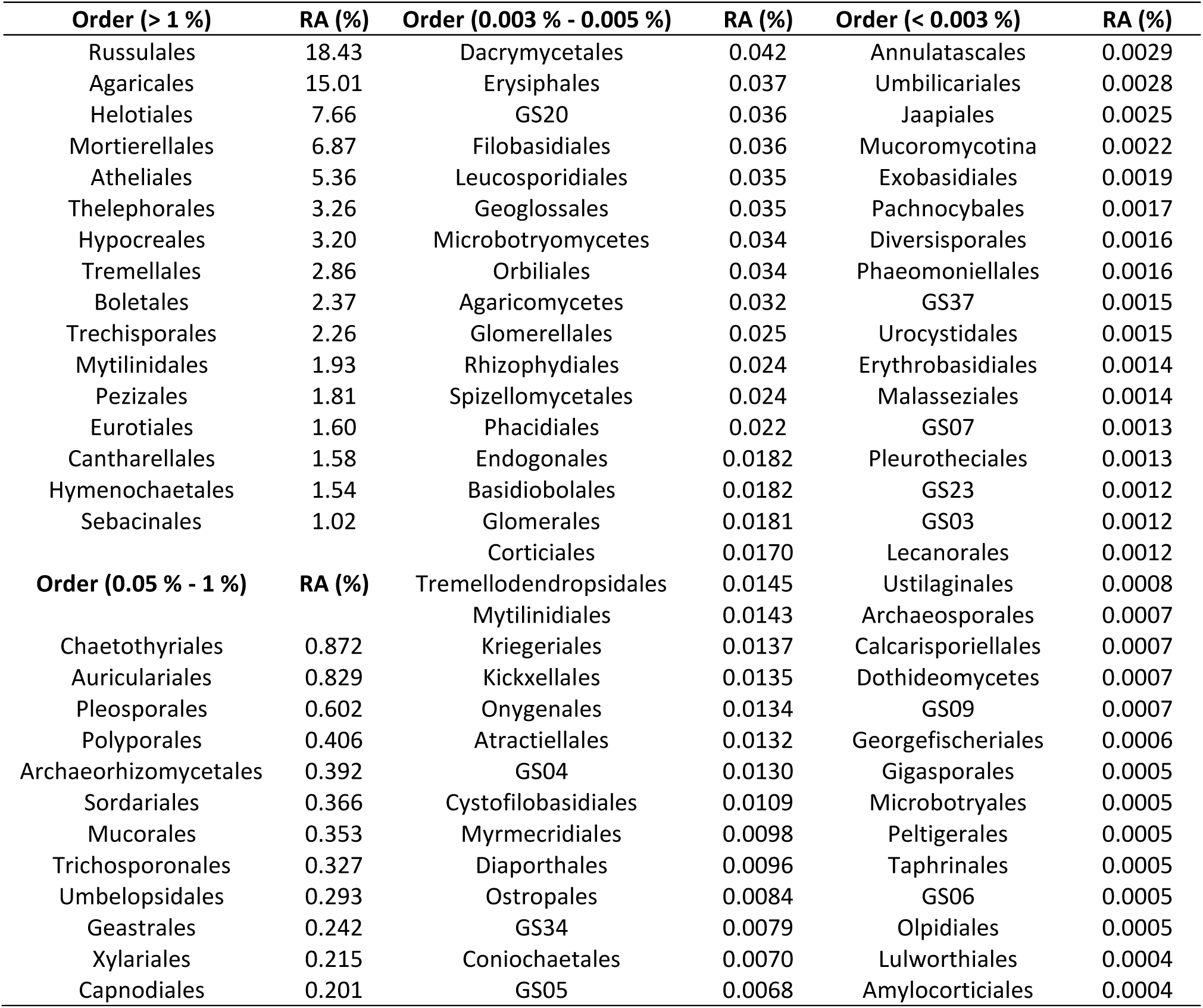
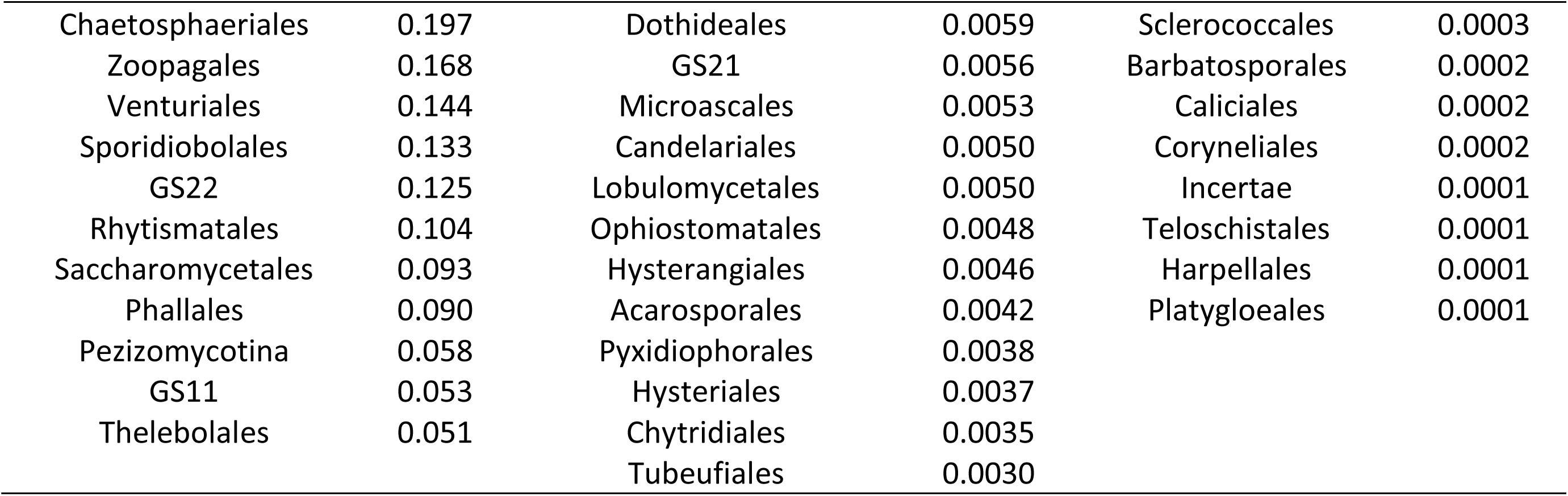
Relative abundance of soil fungal orders in pure (European beech; Douglas-fir; Norway spruce) and mixed (European beech with Douglas-fir; European beech with Norway spruce) tree stands. Relative abundance compiled out of total abundance (1,640,160 reads) comprising all orders.

**Supplement Fig. S1.**
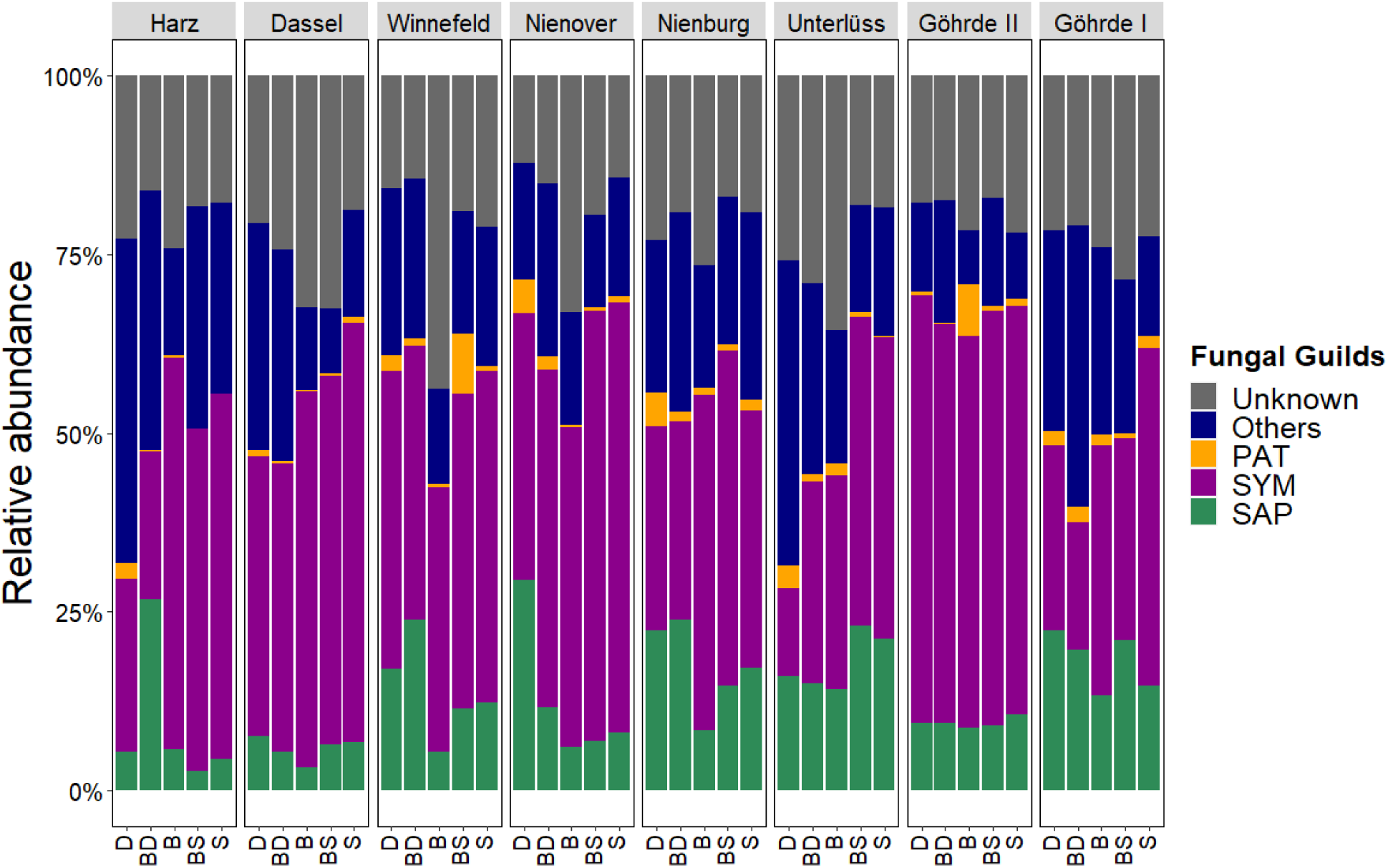
Relative abundance of symbiotrophic (SYM), saprotrophic (SAP), pathotrophic (PAT), others and unidentified (unknown) fungi in soil of the study sites Harz, Dassel, Winnefeld, Nienover, Nienburg, Unterlüss, Göhrde II and Göhrde I in different tree stands of Douglas-fir (D), beech (B), spruce (S), mixture of beech with Douglas-fir (BD) and mixture of beech with spruce (BS). Other fungi are all fungi for which no clear guild annotation was obtained. Unknown fungi lack phylogenetic information. Data show means (n = 3 per site and stand).

**Supplement Fig. S2.**
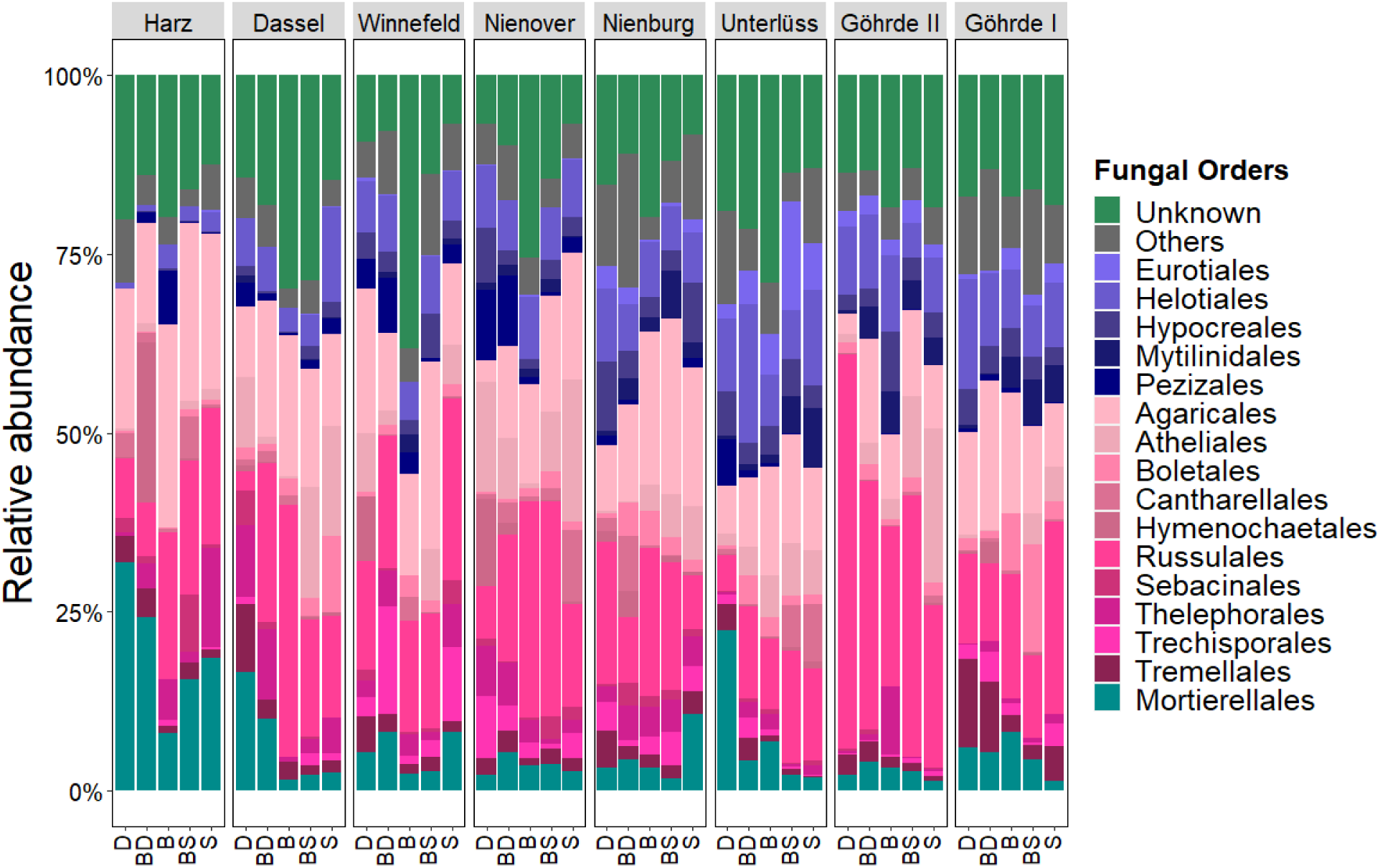
Relative abundance of abundant fungal orders (>1 %) among the different study sites of Harz, Dassel, Winnefeld, Nienover, Nienburg, Unterlüss, Göhrde II and Göhrde I in different tree stands of Douglas-fir (D), beech (B), spruce (S), mixture of beech with Douglas-fir (BD) and mixture of beech with spruce (BS). Orders are arranged according to phyla Ascomycota (blue), Basidiomycota (pink to violet) and Mortierellomycota (dark cyan). Other fungi represent the sum of all fungi with abundances < 1% per order. Unknown fungi lack phylogenetic information. Data show means (n = 3 per site and stand).

**Supplement Fig S3.**
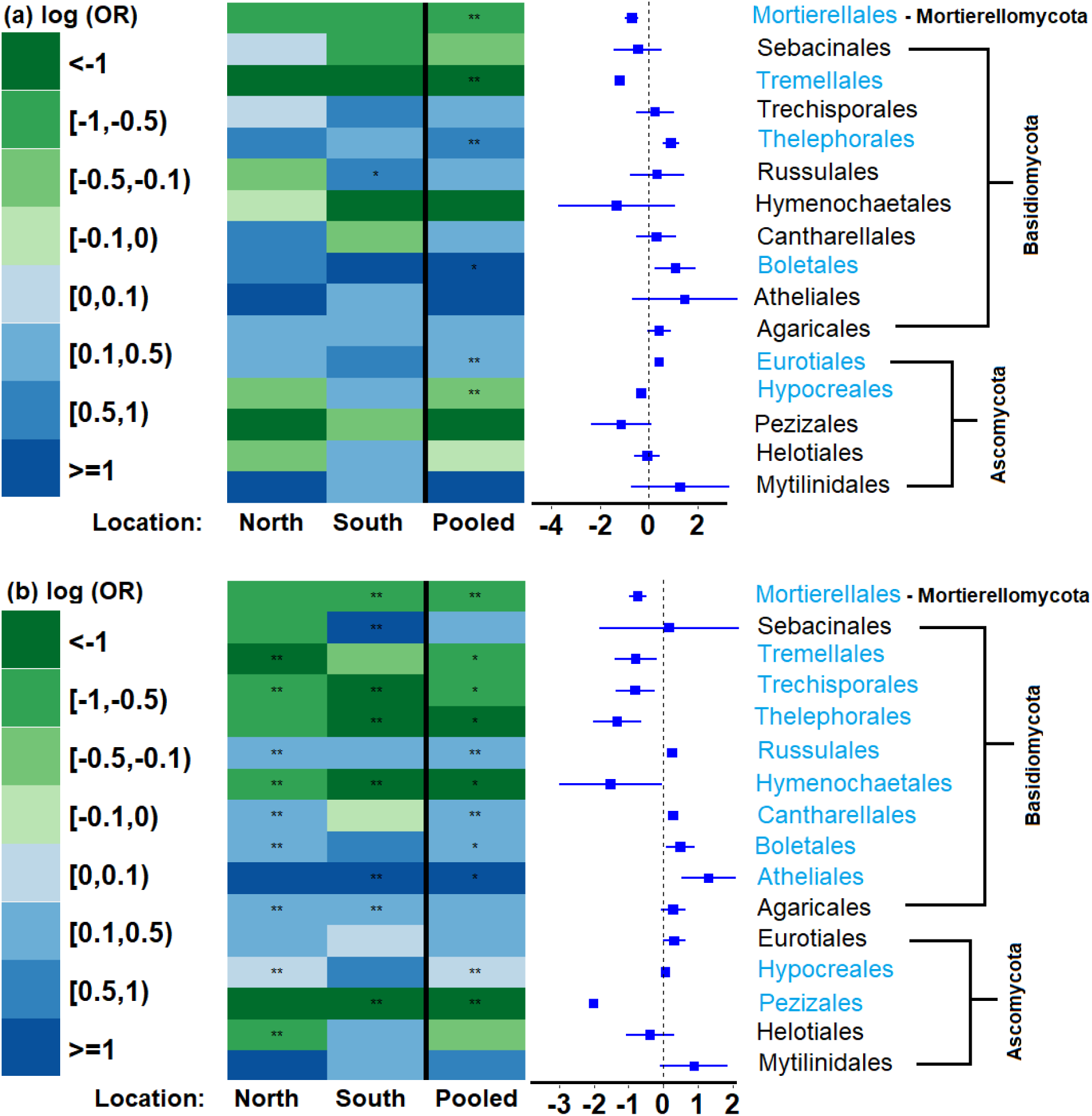
Changes in the relative abundance of fungal orders in conifer and mixed stands: (a) spruce/Douglas-fir, (b) beech-spruce/beech-Douglas-fir. Data are shown as log ratio of odds (log(OR)). Green cells (negative values) indicate enrichment of a fungal order in Douglas-fir and beech-Douglas-fir forests and blue (positive values) indicate enrichment in spruce and beech-spruce forests. Significant differences at p ≤ 0.05 (GAMLSS model with BEZI family, n = 4 for the fungal orders in stands in the north and south location, n = 8 for the pooled data) are indicated with stars (* p < 0.05, ** p < 0.01). Significant orders are highlighted in blue.

